# Banded mongooses discriminate relatedness and MHC diversity in unfamiliar conspecifics

**DOI:** 10.1101/2024.12.22.629965

**Authors:** Nadine Schubert, Carolin Stober, Maibrit Born, Francis Mwanguhya, Robert Businge, Solomon Kyambulima, Kenneth Mwesige, Michael A. Cant, Hazel J. Nichols, Jamie C. Winternitz

## Abstract

Olfactory cues are critical in mammalian social communication, conveying fitness-relevant information such as relatedness, genetic quality, and compatibility. Recognizing kin through scent can help avoid inbreeding depression and guide nepotistic behaviors, enhancing both direct and indirect fitness. While many species use familiarity to identify relatives, others rely on phenotype matching, where animals assess genetic similarity by comparing their own genetically determined odor with that of others. In banded mongooses, synchronized breeding disrupts familiarity cues, increasing reliance on alternative mechanisms for kin discrimination and mate selection. We tested whether banded mongooses use odors to assess genetic diversity and relatedness based on (1) major histocompatibility complex (MHC) genotypes and (2) neutral microsatellite loci that reflect genetic diversity and relatedness. We found that individuals respond differently to odors from unfamiliar individuals based on MHC diversity and genetic relatedness. Specifically, individuals show more interest in less MHC diverse and less related unfamiliar conspecifics, suggesting odor cues are used to evaluate threat level of intruders or competitors. Genetic diversity had no impact on responses to odors and was not significantly associated with MHC diversity, implying that responses to MHC diversity did not result from an underlying correlation with over-all genetic diversity. We also found no effect of MHC similarity, which might be caused by the limited sample size for this analysis. Our findings show that MHC diversity might signal the genetic quality of individuals, but regions of the genome other than MHC may be used to assess relatedness. These findings provide the basis for future research on the involvement of the MHC and other genes in social communication in species where phenotype matching is likely to be advantageous.

## INTRODUCTION

Scent marks in mammals play a key role in social communication, conveying fitness-relevant information such as relatedness, genetic quality, and compatibility (e.g. Charpentier et al. 2008; Stoffel et al. 2015). Recognizing related individuals through olfactory cues helps animals avoid inbreeding depression (Pusey and Wolf 1996), especially in populations where delayed dispersal increases the risk of mating with close relatives (Koenig and Dickinson 2004; Russell 2009; reviewed in Nichols 2017). Additionally, these cues can guide nepotistic behaviors, supporting kin and enhancing indirect fitness (Hamilton 1964). Scent marks may also signal genetic compatibility, influencing mate selection and improving offspring viability (Penn 2002).

Odors reflecting genetic information are particularly intriguing because their nature suggests that they must have a genetic basis. One prominent system for such signaling is the major histocompatibility complex (MHC), which is both highly polymorphic and critical to immune function (Klein 1986; Bjorkman et al. 1987). MHC molecules bind peptides for self/non-self recognition and initiate immune responses against pathogens. MHC class I (MHC-I) molecules, present on nearly all nucleated cells, detect intracellular peptides, while MHC class II (MHC-II) molecules, found on professional antigen-presenting cells, bind extracellular peptides that have been ingested by the APC (Klein 1986; Neefjes et al. 2011). Pathogen-driven evolution of the MHC underpins its extraordinary allelic diversity.

Three main mechanisms—heterozygote advantage, rare allele advantage, and fluctuating selection—are thought to maintain MHC diversity (Spurgin and Richardson 2010; Radwan et al. 2020). Heterozygote advantage allows individuals with more diverse MHC alleles to bind a wider variety of peptides, enhancing pathogen defense (Pierini and Lenz 2018). Rare allele advantage arises because pathogens adapt to common alleles, making rare alleles more effective (Lenz 2018). The MHC’s role in immunity also extends to mate choice, where studies across vertebrates indicate a preference for MHC-diverse partners, potentially enhancing offspring fitness (Kamiya et al. 2014; Winternitz et al. 2017; Winternitz and Abbate 2022). Despite these findings, the mechanisms linking MHC diversity to mate choice and fitness remain poorly understood.

MHC-related information might be transmitted through odor in several ways: (1) MHC molecules shed from cells (Boehm and Zufall 2006), (2) peptides bound by MHC molecules (e.g. Milinski et al. 2005; Spehr et al. 2006; Hinz et al. 2013), or (3) MHC-regulated changes in the microbial community (Singh et al. 1990) or metabolic pathway (Aksenov et al. 2012) that produce odorants (reviewed in Schubert et al. 2021; and Milinski 2022). Animals may use these cues to assess kinship through two main strategies: familiarity or phenotype matching (Lacy and Sherman 1983; Moore and Ali 1984; Todrank and Heth 2003). Familiarity relies on early-life associations to avoid mating with close kin, such as parents or siblings (Berger and Cunningham 1987). Phenotype matching, on the other hand, allows individuals to compare their own scent with that of others to estimate genetic similarity, enabling kin recognition even among unfamiliar individuals (Lacy and Sherman 1983; Todrank and Heth 2003). This flexibility highlights the potential for MHC-mediated cues to influence social and reproductive behavior in diverse contexts.

Banded mongooses are cooperative breeders that show limited dispersal and usually reproduce within their natal pack (Cant et al. 2016). Inbreeding thus occurs frequently and in the population observed for more than 25 years in the Queen Elizabeth National Park in Uganda, two thirds of the population are to some extent inbred, including 7.1% with inbreeding coefficients above 0.25, which results from full-sibling or parent-offspring matings (Wells et al. 2018). Despite inbreeding being widespread, it has been observed to incur a cost on individual fitness in the form of yearling body mass and male reproductive success (Wells et al. 2018). Female banded mongooses can synchronize their estrus resulting in all breeding females giving birth on the same day and combining their pups into a single communal litter (Cant et al. 2016). This breeding behavior likely disrupts familiarity cues (Marshall et al. 2021). However, banded mongooses choose mates that are less closely related than what would be expected by chance (Sanderson et al. 2015) and this pattern cannot be explained by the use of familiarity cues (Khera et al. 2021). Furthermore, banded mongooses appear to discriminate relatedness when evicting members from the group. Females are more likely to evict females that are younger, and they also appear to apply negative kin discrimination, as more closely related females are more likely to be targeted (Thompson et al. 2017). Another context in which discrimination of relatedness may be used is escorting. The synchronized reproduction leads to a communal litter and once the pups leave the den and join the group for foraging trips at approximately 3-4 weeks of age, adults provide pups with food (Cant et al. 2016). This pup-escort relationship is beneficial for the pup, as it increases survival, body weight and faster reproductive onset (Hodge 2005). Although there is no evidence that pup-escort pairs are formed based on relatedness, male escorts have been observed to increase their care by spending more time escorting the pup with growing relatedness between them (Vitikainen et al. 2017).

Mitchell et al. (2018) provided evidence that banded mongooses can discriminate relatedness in familiar individuals despite disrupted familiarity cues. They found interest in odors decreases with increasing relatedness and supposed that phenotype matching might be used to obtain reliable information for mate choice decisions since familiarity should be no useful cue in the case of banded mongooses. However, whichever mechanism is used to discriminate relatedness, it does not seem to be applied in unfamiliar conspecifics (Mitchell et al. 2018). Beyond kin discrimination for inbreeding avoidance, phenotype matching may be used to assess relatedness levels in an intra-sexual selection context as well. One of these contexts is the eviction described in the previous paragraph, during which females negatively discriminate relatedness when the evictees are older (Thompson et al. 2017). For males, assessing genetic quality of potential intruders might be particularly important. Compared to females, males have a greater role in territory defense (Cant et al. 2016) and subordinate males respond first to an intruder, are more aggressive towards them than dominant males and spend more time inspecting them (Cant et al. 2002). Gaining information about the genetic makeup of a potential intruder might aid in assessing the potential threat of the intruder or its potential for competing for matings with the defender and help allocate resources effectively.

Our study aims to investigate whether banded mongooses use odor cues, potentially stemming from the MHC, for discriminating unfamiliar conspecifics. We predicted that banded mongooses i) are more interested in mongoose odors than blank controls, ii) respond to relatedness and MHC diversity in unfamiliar individuals, iii) are less interested in MHC similar odors.

## MATERIAL AND METHODS

### 2.1 Study site

Data used in this study were collected from a wild population of banded mongooses in Queen Elizabeth National Park in Uganda (0°12’S, 27°54E’). The study area consists of approximately 10 km² savannah and includes the Mweya peninsula and the surrounding mainland area. Behavioral, life-history and genetic data as well as information on group composition and territorial structures have been collected regularly and systematically for over 25-years. The population consists of 10-12 packs at any one time, corresponding to approximately 250 individuals. Individuals are identifiable in the field by sight based on (1) dye patterns in the fur that were applied using commercial hair dye (L’Oreal, UK) for individuals up to 6 months old, and (2) shaved fur patterns or (3) color-coded plastic collars for adults that had stopped growing. Shave patterns and collars were maintained during trapping events that took place every 3-6 months as described by Cant (2000), Hodge (2007), and Jordan et al. (2010). Upon first capture, individuals were given either an individual tattoo or a subcutaneous pit tag under anesthetic (TAG-P-122IJ, Wyre Micro Design Ltd., UK) to allow permanent identification, and a 2mm tissue sample was taken from the tip of the tail for genetic analysis.

### 2.2 Odor collection

Banded mongooses scent mark frequently and use it for communication between packs, for example to mark their territory (Jordan 2009), and also to convey information within packs regarding reproductive state (Mitchell, Cant, and Nichols 2017), and relatedness (Mitchell et al. 2018). Thus, we used anal gland secretion (AGS) as the source of odor. We collected AGS from two social groups that were inhabiting non-neighboring territories and thus were unfamiliar with each other.

Samples were collected between May and July of 2022 from 39 adults (≥12 months of age), comprising 9 females and 30 males. These individuals represent all adult individuals from one group and five males from another group. The second group was just recently formed through a fusion of males from a habituated group and unhabituated females, which is why no presentations to those females was possible. These circumstances together with the higher longevity of males (Cant et al. 2016) caused a strong male bias in our sample. Animals were trapped according to the protocol described in Jordan et al. (2011). In short, animals were trapped using Tomahawk traps equipped with bait and anaesthetized using isoflurane. Before extraction of AGS, the skin surrounding the exit of the gland was cleaned using clean cotton wool and Nitrile gloves were worn by the handler during the procedure.

Without touching the gloves, the AGS was then collected in 2.5 ml screw-cap glass vials. Distribution between sample vials was solely performed using glass pipettes or metal spoons to avoid altering the odor. The AGS was then immediately frozen in liquid nitrogen until further usage. Groups used for scent presentations were not in estrus and females were not pregnant during sample collection.

### 2.3 Odor presentations

A total of 336 odor presentations (298 experimental and 38 control) were carried out. Odor samples were removed from the liquid nitrogen and were put on ice in a thermos flask (for a maximum of 90 minutes) until usage in the field. Once a pack was located and individuals resumed foraging, the sample was defrosted and applied to a clean tile using a glass stick or metal spatula. The tile was then presented to the focal individual while it was foraging at least 1 m away from other conspecifics, as described in Mitchell et al. (2017). The individual’s response was filmed using a handheld camera. The tile was cleaned with hot water and baking soda using a brush after every presentation. A negative control was conducted for each individual to make sure individuals were not responding to the novelty of the tile itself, but to the odor presented. For this reason, individuals were presented with a clean tile that contained no odor sample. Each individual was only presented one odor per day and after two days of presentations the pack was given one day without presentations.

Moreover, if a presentation was interrupted, e.g. an individual inspecting the odor was startled by a warning call or pushed away from the odor by another individual, repetition of the presentation of this odor was shifted as far to the end of the field season as possible. Both measures were implemented to avoid habituation to the odor and thus changes in the response to it.

### 2.4 Video analysis of responses

Videos were analyzed independently by two people using BORIS software (Friard and Gamba 2016). Responses to the odor presented were categorized as (i) time spent in proximity (one body-tail length) to the odor (= duration) (ii) time spent directly above or touching the tile containing the odor (= contact), and (iii) marking behaviors. Duration and contact both started once the nose of the individual was above the tile. Contact time was measured until the individual either stopped touching the tile with a body part or until it stopped holding its head above the tile. ‘Duration time’ continued until the individual resumed foraging, laid down, groomed other individuals, or moved away from the tile with a distance of at least one body-tail length. Contact behaviors could be split further into sniffing, licking and rolling. Marking behavior included overmarking as well as markings in the vicinity of the tile (one body-tail length) and could either be urine, feces or AGS markings. As concluded by Mitchell et al. (2017), these measures are not independent of one another, as an increased number of vicinity marks increases the duration spent in the vicinity of the odor and thus need to be interpreted accordingly. Moreover, since we didn’t know in which context MHC diversity might influence behavior, we included both overmarking, which may have a competitive function (Rich and Hurst 1999; Wolff et al. 2002; Jordan et al. 2011), as well as vicinity marks, which are thought to be important in mate-choice decisions (Rich and Hurst 1999), in the analysis.

### 2.5 Neutral genetic analyses

We extracted DNA from 39 individuals using the Qiagen® DNeasy blood and tissue kit according to the manufacturers protocol. These individuals were genotyped at 35-43 neutral microsatellite loci based on the methods described in detail by Sanderson et al. (2015). Individual standardized multilocus heterozygosity (sMLH) was calculated using the R package InbreedR (Stoffel et al. 2016). Genetic marker-based relatedness (Queller and Goodnight 1989) was estimated using GENALEX (Peakall and Smouse 2006). One individual was removed from the analysis as it had only been genotyped at 5 microsatellite loci, so relatedness and heterozygosity estimates were potentially unreliable (all other individuals had been genotyped at a minimum of 18 loci).

### 2.6 MHC genetic analyses

MHC genotyping was carried out using target enrichment and PacBio long-read sequencing described in Winternitz, J.C., Schubert, N., Heitlinger, E., Foster, R. G., Cant, M.A., Mwanguhya, F., Businge, R., Kyambulima, S., Mwesige, K., Nichols H.J. (unpublished data). First, a custom hybridization panel was designed by Twist Bioscience to be compatible with PacBio HiFi long reads. Briefly, 68 banded mongoose MHC sequences (GenBank Accession numbers PQ7681 - PQ7748) were blast searched against the banded mongoose genome NCBI GenBank HiC chromosomal assembly GCA_028533875.1 (Megablast, max e-value = 1e-50, maximum hits=1 per sequence). These exon sequences matched 17 unique scaffolds. We restricted potential hybridization targets to those from scaffolds at least 2000 bp long, leading to 14 genomic regions across 12 scaffolds (3 on scaffold/chromosome 8) that included 9 putative class I loci and 5 putative DRB loci.

Next, hybridization, library preparation and sequencing were carried out at Edinburgh Genomics according to the manufacturers’ protocols. Briefly, 39 banded mongoose samples with average DNA concentration of 6.7 ng/ul (range=0.2-29.0, SD=6.6) were used to create PacBio libraries. After size selection, post-PCR quantification, and malfunction in the PacBio Sequel IIe system, only 27 banded mongoose DNA samples had high enough concentration and quality to create PacBio libraries for sequencing. Hybridization was carried out using version ‘REV 2’ of the Twist library preparation and enrichment protocol and a Twist custom panel of mongoose probes. The final sequencing library was loaded on a PacBio Sequel IIe system and produced 173,486 total HiFi reads. Samples were demultiplexed and PCR duplicate reads were removed prior to downstream processing using pbmarkdup v1.0.3 (https://github.com/PacificBiosciences/pbmarkdup), leaving 127,716 unique reads, mean 4912 (st.d. = 3151) per sample. Genotyping was carried out as described in Winternitz, J.C., Schubert, N., Heitlinger, E., Foster, R. G., Cant, M.A., Mwanguhya, F., Businge, R., Kyambulima, S., Mwesige, K., Nichols H.J. (unpublished data).

Briefly, for each sample, HiFi reads were assembled *de novo* into diploid-aware contigs using hifiasm (Cheng et al. 2021) and blast searched against the custom Twist target panel. Contigs of interest were then aligned using MAFFT (Katoh et al. 2002; Katoh and Standley 2013) and a maximum likelihood phylogenetic tree was created using IQTree (Nguyen et al. 2015; Kalyaanamoorthy et al. 2017) to identify monophyletic putative loci. Consensus reference loci sequences were created using custom R code and for each individual raw HiFi reads were mapped to these references using pbmm2 v1.0.3 (https://github.com/PacificBiosciences/pbmarkdup). Variants were called using DeepVariant (Poplin et al. 2018) and haplotypes phased using WhatsHap (Martin et al. 2016). Consensus reference loci were annotated using carnivore NCBI reference sequence MHC annotations and Exonerate v. 2.4.0 (Slater and Birney 2005) and these gene annotations were transferred to individuals’ haplotypes using liftoff (Shumate and Salzberg 2021). In total, individuals were genotyped at 7 MHC-I genes and 7 MHC-II genes. As the number of alleles per individual increased with HiFi read count (Pearson’s cor=0.560, p-value = 1.637e-10), the number of unique HiFi reads was included in downstream analyses.

Allele and supertype sharing were calculated to estimate MHC similarity between individuals. Allele and supertype sharing between individuals were calculated as twice the sum of alleles the individuals shared divided by the sum of alleles of both individuals: D=2Fab/(Fa+Fb) (Wetton et al. 1987). MHC and supertype diversity were estimated as the total number of alleles and supertypes in an individual. Supertypes were estimated using amino acid distances between sequences and then grouped based on functional similarity. The Sandberg distance (Sandberg et al. 1998), 5 physio-chemical z-descriptor values, was calculated for each MHC peptide binding residue (MHC-I (Saper et al. 1991); MHC-II (Brown et al. 1993)) using the R package ‘Peptides’ (Osorio et al. 2015), and transcribed into a similarity matrix. To these matrices we applied find.cluster() using the criterion “goesup” and method=”kmeans” for MHC-I and criterion “diffNgroup” and method=”ward” for MHC-II. This method was repeated 1000 times and the mean, mode, and median number of clusters calculated to arrive at 11 and 10 clusters, respectively. We assigned alleles to groups using the dapc() function (i.e., discriminant analysis of principal components) from the R package ‘adegenet’ (Jombart 2008) and repeated this process 1000 times to estimate repeatability with light’s kappa value in ‘irr’ R package (Gamer et al. 2019) For MHC-I, repeatability Kappa = 1 and the mean assignment proportion was 0.988. For MHC-II repeatability was perfect, with Kappa = 1 and the mean assignment proportion = 1.

### 2.7 Statistical analysis

#### 2.7.1 Correlational analysis

Strong collinearity among variables included in statistical models can impede model interpretation (reviewed in Harrison et al. 2018). Therefore we used Pearson’s product-moment correlation from R version 4.4.0 (R Core Team 2023) to investigate the degree of correlation between (1) MHC diversity (the number distinct alleles per individual) and genomic diversity (sMLH), (2) MHC allele similarity and relatedness, and (3) MHC supertype similarity and relatedness. Since response measures are not independent of one another, as the time spent sniffing an odor or marking should correlate with the time spent in proximity to an odor (Mitchell, Cant, and Nichols 2017), we also investigated potential collinearities between all behavioral response variables (Contact, Sniffing, Duration, Licking, Marking and Rolling).

#### 2.7.2 General linear mixed models

First, to establish that banded mongooses respond to anal gland odors, rather than to a novel object in their environment, we tested for a difference between control and experimental presentations using six general linear mixed effect models (GLMMs). Detailed methods for these correlational analyses can be found in the supplementary materials.

Second, we investigated whether banded mongooses responded to the sex and genetic diversity of the odor donor, and to the relatedness between the donor and recipient. MHC measures were not included in this model to maximize the size of the dataset (only a subset of individuals had MHC data available for them). In three separate GLMMs, we modeled our response variables (contact (log), sniffing (log +1) or duration (log)) predicted by the sex of the donor and recipient, genetic relatedness and sMLH. The identity of the odor donor and recipient were included as random effects. The packs of the odor donor and recipients were not included as random effects because the variance explained by them was low, including them usually led to a singular fit, and the effect of the pack should be subsumed within the individual identities of the pack members. This analysis included 308 odor presentations: male to male (N=230), male to female (N=43), and female to male (N=35). No female odor was presented to females, as the females of one of the packs were not sufficiently habituated to perform presentations. Odor presentations involved 37 individual recipients and 35 individual odor donors.

Third, to investigate the effect of MHC diversity on the animals’ response, we fitted three GLMMs, each with one of our response variables (contact (log), sniffing (log +1) or duration (log)). The number of unique MHC alleles was included as an explanatory variable, along with sMLH (to control for background genomic diversity), the number of HiFi reads (to control for sequencing effort) and the sex of the odor donor and recipient. The identity of the odor recipient was included as a random effect. The identity of the odor donor was not included as a random effect due to it explaining zero variance and causing a singular fit, likely because the variance associated with the odor donor was related to its sex. These models included 111 experimental odor presentations from 17 donors to 36 individuals (male to female: N=9, female to male: N=20, male to male: N = 82, no female odor was presented to females).

Finally, we investigated whether banded mongooses responded differently to odors based on MHC similarity between odor donor and recipient. Since MHC similarity data requires MHC genotypes to be available for both odor donor and recipient, there was a very limited number of data points (N=33 presentations to 19 individuals – 13 males and 6 females; male to female: N = 6, female to male: N = 4, male to male: N = 23) available for this analysis. As with previous analyses, we fitted three models, one for each response variable (contact (log), sniffing (log +1) and duration (log)). MHC similarity between donor and recipient was fitted as the predictor variable, and recipient ID was fitted as a random effect. Due to the small dataset, the model was reduced to a single predictor and random term to retain sufficient statistical power and avoid overfitting.

All GLMMs were constructed in R version 4.4.0 (R Core Team 2023) using the lme4 package (Bates et al. 2017) and were fitted with a Gaussian family. Significant fixed effects were detected using the R package *afex* version 1.4-1 (Singmann et al. 2018) with Type III Analysis of Variance with Satterthwaite’s method.

## RESULTS

### 3.1 Preliminary analyses

Correlations between the MHC diversity measures showed highly significant and strong correlations between MHC allele number and supertype number (r = 0.883, p < 2.2^-16^) and MHC allele similarity and MHC supertype similarity (r = 0.835, p = 1.488^-9^). Since both allele number and supertype number contain different levels of information on functional diversity of the MHC, we decided to include all measures in our analyses but in separate models. Correlations between the response measures showed a strong and highly significant correlation between contact and rolling (r = 0.737, p < 2.2^-16^). However, comparison between responses to control and experimental presentations indicated that only contact, sniffing and duration were significantly different and thus only these response measures were considered for further analysis. Contact and duration also correlated significantly (r = 0.698, p < 2.2^-16^), but we decided to include both in the analysis, as they might contain different behaviorally relevant information, and the variables were also fitted in separate models. Detailed results of the correlational analyses and the comparison of control and experimental presentations can be found in the supplementary material (Tab. S1 & S2).

### 3.2 Influence of sex, relatedness and sMLH

Banded mongooses responded differently to odors dependent on the sex of the odor donor, the sex of the recipient, and the genetic relatedness between the odor donor and recipient (Tab. 1). However, the genetic diversity of the donor did not impact behavioral responses of the recipient. Specifically, males spent more time in contact with the presentations, sniffing them, and had longer behavioral responses in comparison to females, and responses towards female odors were elevated compared to responses towards males for contact and duration (Fig. 1). Finally, the duration of the response to odors decreased as relatedness between the donor and recipient increased (Fig. 2).

**Figure 1.**
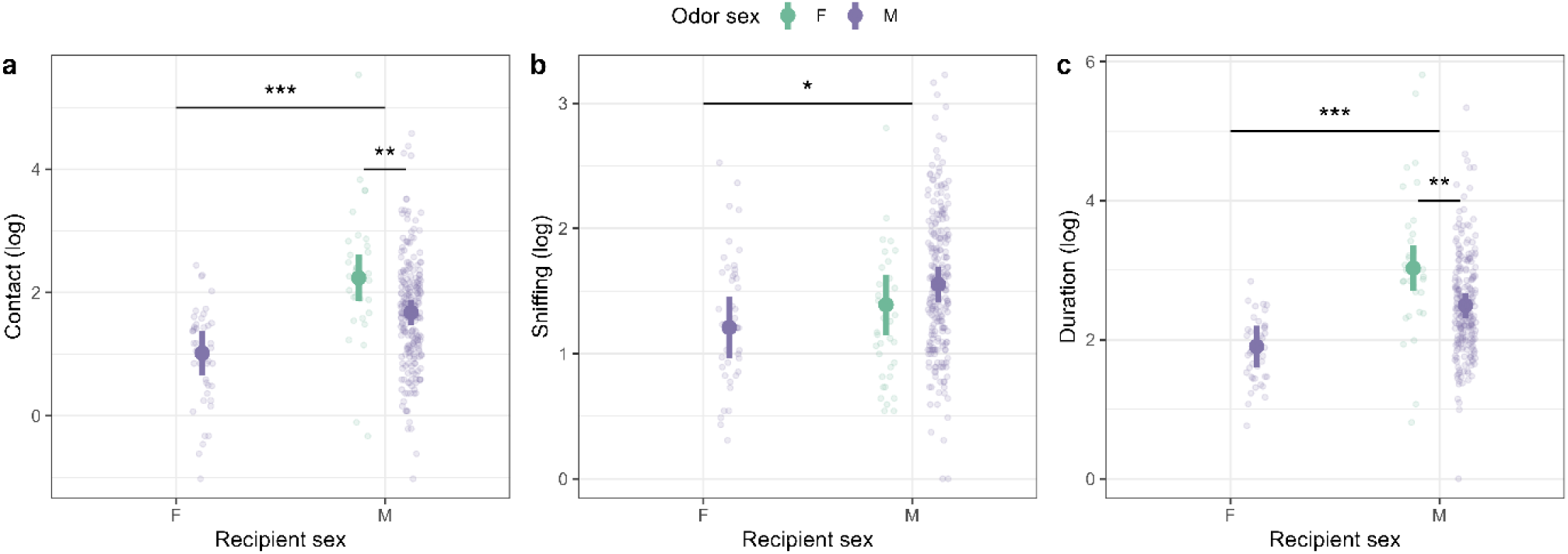
Sex-dependent responses. Predicted model responses towards male and female odors measured as contact (a), sniffing (b) or duration (c) time shown separately for the sexes and colored by the sex of the odor donor.

**Figure 2.**
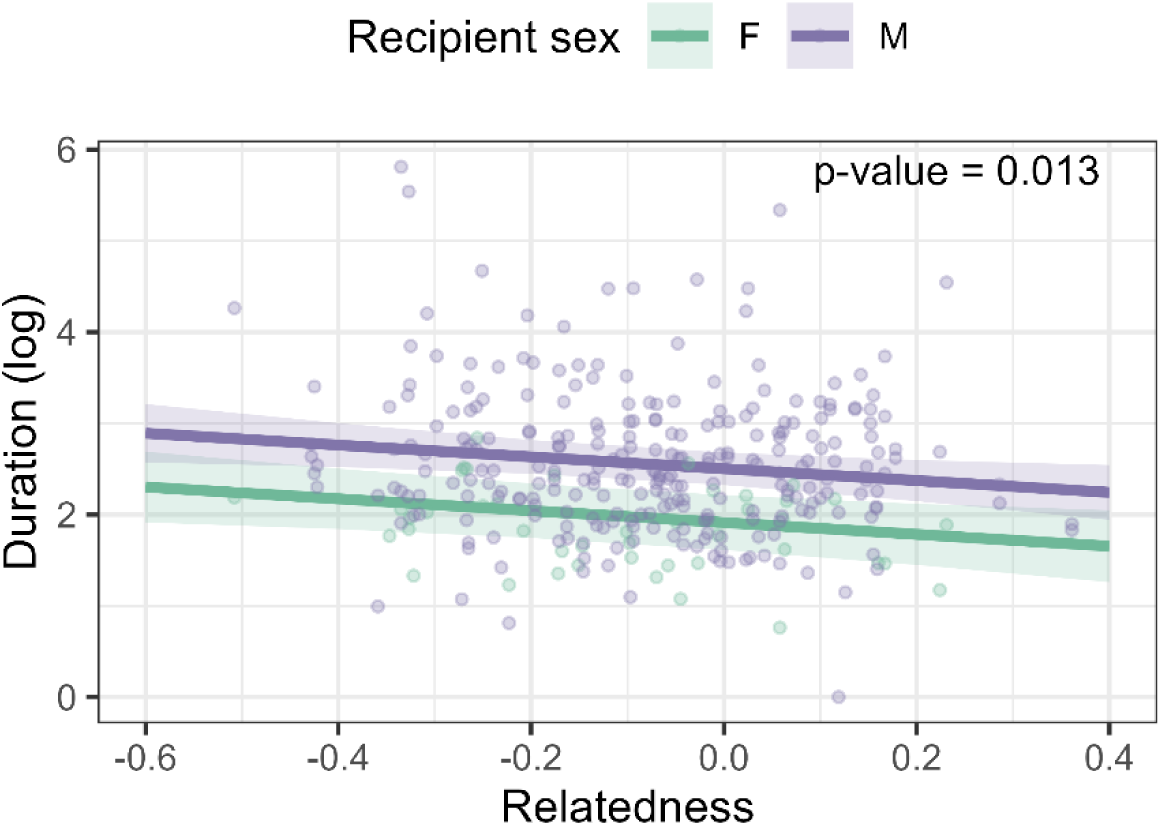
Duration response declines with increasing relatedness. Responses are model predictions controlling for other predictor variables and for repeated measures of recipient ID and donor ID as random effects. Raw data are superimposed for visualization. Relatedness values are negative if the value of an individual is below that of the average in the population.

**Table 1.** Model output for models investigating influences affecting responses. Model output for effects of genetic relatedness between recipient and odor donor on contact, sniffing and duration responses. P-values were calculated based on Satterthwaite’s method.

| Response variable | Fixed effect | estimate | SE | p-value |
| --- | --- | --- | --- | --- |
| Contact | Recipient sex | 0.6550 | 0.1831 | **0.00073\*** |
|  | Odor sex | -0.5594 | 0.1782 | **0.0029\*** |
|  | sMLH | 0.3116 | 0.4915 | 0.5325 |
|  | relatedness | -0.3376 | 0.3177 | 0.2887 |
| Sniffing | Recipient sex | 0.3392 | 0.1272 | **0.0107\*** |
|  | Odor sex | 0.1650 | 0.1083 | 0.1325 |
|  | sMLH | -0.0205 | 0.3008 | 0.7477 |
|  | relatedness | 0.0623 | 0.1934 | 0.9460 |
| Duration | Recipient sex | 0.5894 | 0.1397 | **7.12<sup>-05\*</sup>** |
|  | Odor sex | -0.5352 | 0.1650 | **0.0025\*** |
|  | sMLH | 0.1569 | 0.4712 | 0.7419 |
|  | relatedness | -0.6490 | 0.2609 | **0.0134\*** |

### 3.3 Influence of MHC diversity

MHC diversity of the odor donor measured as the number of distinct alleles had no significant effect on contact and sniffing, and duration was borderline insignificant and negatively linked to MHC diversity (estimate = −0.04678, SE = 0.0236, p= 0.054, Fig. 3a, Tab. 2) indicating that time spent in the vicinity of the odor before resuming foraging or resting behaviors decreased with increasing MHC diversity. Male recipients still spent more time close to the odor and in the vicinity of the odor than female recipients in these smaller dstasets (contact: estimate = 0.8461, SE = 0.3563, p = 0.0201, duration: estimate = 0.9295, SE = 0.2863, p = 0.0017).

**Figure 3.**
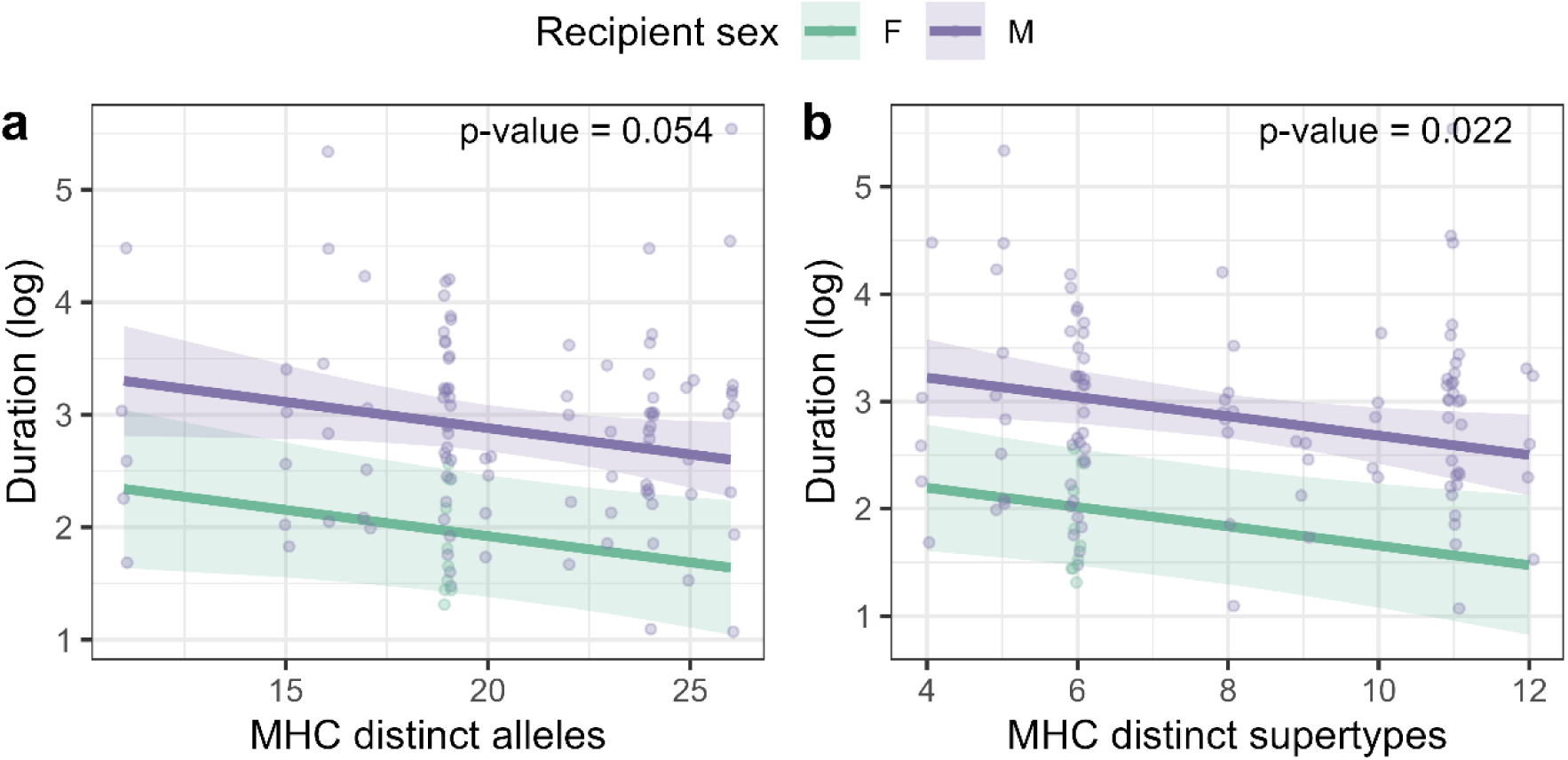
Duration response in relation to MHC diversity. Duration response in relation to MHC distinct alleles (a) and supertypes (b). Responses are model predictions controlling for the effect of sex.

**Table 2.** Model outputs for relationships with MHC diversity. Model output for effects of MHC diversity of the odor donor measured as the number of distinct alleles or supertypes respectively on contact, sniffing and duration responses.

| MHC measure | Fixed effect | Response variable | estimate | SE | p-value |
| --- | --- | --- | --- | --- | --- |
| Alleles | Recipient sex | Contact | 0.8461 | 0.3563 | **0.0201\*** |
|  | Odor sex |  | -0.2689 | 0.3186 | 0.4008 |
|  | sMLH |  | -0.2478 | 0.8396 | 0.7686 |
|  | MHC diversity |  | -0.0142 | 0.0281 | 0.6151 |
|  | Unique reads |  | 5.915<sup>-05</sup> | 5.823<sup>-05</sup> | 0.3122 |
|  | Recipient sex | Sniffing | 0.3304 | 0.2430 | 0.1819 |
|  | Odor sex |  | 0.1240 | 0.1985 | 0.5344 |
|  | sMLH |  | 3.043<sup>-03</sup> | 0.5259 | 0.9954 |
|  | MHC diversity |  | 0.0189 | 0.0175 | 0.2837 |
|  | Unique reads |  | -2.051<sup>-05</sup> | 3.652<sup>-05</sup> | 0.5761 |
|  | Recipient sex | Duration | 0.9295 | 0.2863 | **0.0017\*** |
|  | Odor sex |  | -0.2797 | 0.2678 | 0.2990 |
|  | sMLH |  | -0.3284 | 0.7031 | 0.6415 |
|  | MHC diversity |  | -0.04678 | 0.0236 | 0.0506 |
|  | Unique reads |  | 8.877<sup>-05</sup> | 4.871<sup>-05</sup> | 0.0714 |
| *Supertype* | *Recipient sex* | *Contact* | 0.8681 | 0.3572 | **0.0174\*** |
|  | *Odor sex* |  | -0.2633 | 0.3177 | 0.4094 |
|  | *sMLH* |  | -0.1158 | 0.8713 | 0.8946 |
|  | *Distinct supertypes* |  | -0.0330 | 0.0460 | 0.4707 |
|  | *Unique reads* |  | 6.728<sup>-05</sup> | 5.990<sup>-05</sup> | 0.2643 |
|  | *Recipient sex* | *Sniffing* | 0.3267 | 0.2443 | 0.1891 |
|  | *Odor sex* |  | 0.1148 | 0.1996 | 0.5672 |
|  | *sMLH* |  | 0.0717 | 0.5486 | 0.8886 |
|  | *MHC diversity* |  | 1.168<sup>-03</sup> | 0.0290 | 0.9860 |
|  | *Unique reads* |  | -5.336<sup>-06</sup> | 3.771<sup>-05</sup> | 0.8879 |
|  | *Recipient sex* | *Duration* | 0.9964 | 0.2860 | **0.0008\*** |
|  | *Odor sex* |  | -0.2606 | 0.2658 | 0.3294 |
|  | *sMLH* |  | -7.563<sup>-03</sup> | 0.7277 | 0.9917 |
|  | *MHC diversity* |  | -0.0879 | 0.0383 | **0.0240\*** |
|  | *Unique reads* |  | 1.033<sup>-04</sup> | 5.004<sup>-05</sup> | **0.0420\*** |

The number of distinct supertypes per individual, representing the amount of functional diversity of an individual rather than allelic diversity, showed a significant negative relationship for the duration response (Tab. 2; estimate = −0.0879, SE = 0.0383, p = 0.0240), controlling for the number of HiFi reads (Fig. 3b). This indicates that individuals spent more time in the vicinity of odors that had fewer supertypes before resuming foraging or resting behaviors. In this model we again found that male recipients had higher contact responses (estimate = 0.8681, SE = 0.3572, p = 0.0174) and longer duration spent near the odor (estimate = 0.9964, SE = 0.2860, p = 0.0008).

### 3.4 Influence of MHC similarity

Neither MHC similarity of the alleles nor of the supertypes significantly influenced contact, sniffing or duration responses nor any of the other fitted variables (Tab. 3).

**Table 3.** Model output for models on MHC similarity. Model output for effects of MHC similarity between recipient and odor donor on contact, sniffing and duration responses.

| *MHC measure* | *Fixed effect* | *Response variable* | *estimate* | *SE* | *p-value* |
| --- | --- | --- | --- | --- | --- |
| *Alleles* | *MHC similarity* | *Contact* | 0.6880 | 1.364 | 0.6186 |
|  | *Unique reads* |  | 2.872<sup>-05</sup> | 7.798<sup>-05</sup> | 0.7182 |
|  | *MHC similarity* | *Sniffing* | -0.1172 | 1.143 | 0.9192 |
|  | *Unique reads* |  | -2.100<sup>-05</sup> | 6.236<sup>-05</sup> | 0.7416 |
|  | *MHC similarity* | *Duration* | -1.084 | 1.210 | 0.3777 |
|  | *Unique reads* |  | 4.729<sup>-05</sup> | 7.227<sup>-05</sup> | 0.5182 |
| Supertype | MHC similarity | Contact | 1.353 | 1.114 | 0.2360 |
| | Unique reads | | $3.260^{-05}$ | $7.788^{-05}$ | 0.6819 |
|  | MHC similarity | Sniffing | 0.4178 | 0.9487 | 0.6640 |
| | Unique reads | | $-2.129^{-05}$ | $6.295^{-05}$ | 0.7408 |
|  | MHC similarity | Duration | -0.369 | 1.006 | 0.7615 |
| | Unique reads | | $4.154^{-05}$ | $7.275^{-05}$ | 0.5728 |

## DISCUSSION

We found that banded mongooses varied in their responses to unfamiliar odors based on genetic relatedness to the odor donor and MHC diversity but not on MHC similarity or overall genetic diversity. Responses also differed depending on the sex of the odor donor and recipient. These findings provide evidence for discrimination of genetic relatedness and MHC diversity in unfamiliar individuals’ odors, suggesting that banded mongooses may employ kin recognition mechanisms like phenotype matching (Holmes and Sherman 1982; Lacy and Sherman 1983; Hepper 1991) to assess genetic information in conspecifics.

Banded mongooses face a high risk of inbreeding due to limited dispersal, with over 80% of individuals remaining in their natal pack (Cant et al. 2016). In our study population, 64% of pups are born to females mating with resident males (Nichols et al. 2014), resulting in more than 7% of pups being offspring of first-order inbreeding, such as parent-offspring or full-sibling matings (Wells et al. 2018). This inbreeding has significant fitness costs, including increased parasite load (Mitchell, Vitikainen, Wells, et al. 2017), reduced yearling body mass, and lower reproductive success in males (Wells et al. 2018). Identifying kin during mate selection could help mitigate these risks. Supporting this, inbreeding occurs less often than expected by chance, and males preferentially mate-guard less related females (Sanderson et al. 2015). As these patterns cannot be explained by familiarity-based cues (Khera et al. 2021), other mechanisms, such as phenotype matching, may be involved. Banded mongooses also appear to discriminate kin in other contexts, including cooperative behaviors (Vitikainen et al. 2017) and competitive interactions (Thompson et al. 2017).

Mitchell et al. (2018) demonstrated that banded mongooses can differentiate odors based on relatedness among familiar group members, but it was unclear whether this discrimination was due to the odors themselves or prior knowledge of the individuals. They found no evidence of relatedness discrimination in unfamiliar individuals, though their relatively small sample size (N = 121 presentations) may have limited the analysis. In contrast, our study used a larger sample (N = 308 presentations) of unfamiliar individuals and found that relatedness significantly influenced the duration of responses to odors. By exclusively testing unfamiliar individuals, we eliminate the confounding effect of familiarity, providing strong evidence that banded mongooses use odor-based cues to assess relatedness via phenotype matching.

While familiarity is a common proxy for relatedness (Pusey and Wolf 1996), it may be insufficient in species where reproductive and social dynamics complicate the use of associative learning. In cooperative species with high reproductive skew—such as meerkats, where dominant pairs monopolize reproduction (Sharp and Clutton-Brock 2010)—familiarity might suffice for kin discrimination. Interestingly, even in meerkats, phenotype matching via odor has been suggested as a complementary mechanism (Leclaire et al. 2013). For banded mongooses, which exhibit low reproductive skew and highly synchronized breeding among dominant and subordinate individuals (Gilchrist 2006), kin recognition via familiarity is less feasible. Their communal litters, formed by multiple females giving birth simultaneously, often include offspring with different paternities, necessitating a mechanism like phenotype matching to assess relatedness independently of familiarity.

Other cooperative species also demonstrate phenotype matching for kin discrimination. African cichlids use visual and chemical cues to assess relatedness among separately reared individuals (Le Vin et al. 2010), and African clawed frog tadpoles apply MHC-based self-referencing to distinguish kin (Villinger and Waldman 2008). However, disentangling phenotype matching from learned familiarity remains challenging. For example, while baboons exhibit preferential treatment of genetic offspring over unrelated offspring from consorts, it remains unclear whether this is due to genetic recognition or behavioral cues, such as perceived mating effort with the mother (Buchan et al. 2003). Studies must carefully account for these confounding factors, recognizing that familiarity and phenotype matching are not mutually exclusive and may operate in tandem (Porter 1988; Tang-Martinez 2001).

In banded mongooses, existing evidence from other studies suggests that phenotype matching may not provide precise relatedness assessment. This imprecision could explain the persistence of inbreeding (Wells et al. 2018), even though mongooses tend to mate with less closely related individuals compared to random mating (Sanderson et al. 2015). Interestingly, this uncertainty in phenotype matching may also support synchronized breeding, which facilitates cooperative behavior. For instance, while banded mongoose females cannot distinguish their own offspring within communal litters, nor can pups identify their mothers (Marshall et al. 2021), breeding asynchrony can lead to infanticide (Hodge et al. 2011). Non-breeding females, not risking their own offspring, are more likely to commit infanticide, causing litters to fail within the first week (Cant et al. 2014). Conversely, synchrony in breeding results in mixed-parentage litters that rarely fail early, likely because imprecise relatedness cues prevent females from risking harm to their own pups.

This inability to assess relatedness precisely may also facilitate a “veil of ignorance,” which promotes equal contributions in cooperative behaviors such as communal offspring care (Marshall et al. 2021). Such mechanisms are thought to enhance cooperation by minimizing kin discrimination, as seen in other species (reviewed in Queller and Strassmann 2013). For example, social insects could theoretically discriminate between patrilines using self-referencing for phenotype matching but instead use colony-wise phenotypes as a template, preventing patriline-specific discrimination (Keller 1997; Queller and Strassmann 2002). Similarly, male birds can differentiate between broods sired from other males and avoid raising them, yet they do not favor their own offspring within mixed broods (Keller 1997). In banded mongooses, the ability to detect relatedness without consistently applying this information aligns with theoretical predictions that uncertainty in kin recognition can promote cooperative behavior (Frank 2003; Okasha 2012; Queller and Strassmann 2013).

The MHC plays a key role in immune response and has the potential to generate odor cues, directly or indirectly (reviewed in Schubert et al. 2021), providing information about an individual’s genetic makeup. Kin discrimination based on MHC-derived odor cues has been observed in various species and contexts. For instance, house mice exhibit a preference for communal nesting with relatives to reduce infanticide and exploitative risks when caring for pups, using MHC similarity as a cue (Manning et al. 1992). Similarly, African clawed frog tadpoles prefer half-siblings sharing MHC alleles, likely employing a self-referencing mechanism (Villinger and Waldman 2008). In mate selection, animals may make use of MHC-related odor cues to increase offspring MHC diversity (e.g. Schwensow et al. 2008), potentially enhancing genomic diversity and reducing inbreeding risks (Jennions 1997; Tregenza and Wedell 2000; Mays and Hill 2004).

In our study, banded mongooses responded to genetic relatedness in odors but showed no evidence of MHC similarity influencing these responses. Furthermore, MHC similarity did not correlate with genomic relatedness. This could stem from the small sample size (n = 33 presentations) or subtle effects requiring larger datasets to detect (Gaigher et al. 2019). Despite this, mongooses demonstrated altered responses to odors based on the diversity of MHC alleles and supertypes in the donor, independent of overall genomic diversity. These findings suggest direct detection of MHC diversity in odors. Furthermore, males showed less interest in more MHC diverse odors, potentially because more MHC diverse males are stronger potential competitors. In support of this, prior work links male banded mongoose reproductive success to higher MHC diversity (Schubert et al. 2024), highlighting MHC diversity’s relevance in fitness and possibly mate choice contexts, as seen across species (Kamiya et al. 2014).

Relatedly, male banded mongooses responded more strongly to unfamiliar odors than females (Mitchell et al. 2018), consistent with their greater role in territorial defense (Cant et al. 2016). Subordinate males are often the first to confront intruders, displaying increased aggression and inspection behaviors (Cant et al. 2002). Extra-group paternity, accounting for approximately 18% of offspring (Nichols et al. 2015), also offers rare reproductive opportunities for subordinate males during inter-pack encounters (Green et al. 2024). These dynamics suggest males have a dual motivation to assess unfamiliar individuals for sex, genetic quality, and compatibility, as such encounters can impact their fitness both positively and negatively.

Similar to communally nesting mice that use MHC-based odor cues to identify relatives and reduce exploitation risks (Manning et al. 1992), banded mongooses may utilize MHC-related odor differences to evaluate the relatedness and threat level of intruders or competitors. This ability could facilitate decisions in social and reproductive contexts, aligning with broader patterns of MHC-based discrimination observed in other species.

## OUTLOOK

Our study provides first evidence that phenotype matching might be used to discriminate relatedness levels and MHC diversity in unfamiliar conspecifics in banded mongooses. Given the high risk of inbreeding in banded mongoose groups, phenotype matching might have evolved as an inbreeding avoidance mechanism. However, it might also be used to assess information about intruders and potential competitors for matings. Future studies should be planned strategically with genotyping of each individual as the first step to allow for ideal MHC combinations in odor recipient and donor and use sample sizes large enough to allow investigating same- and opposite-sex contexts separately. Habituation-dishabituation trials could additionally help with determining exactly what extent of difference in relatedness and MHC banded mongooses are able to discriminate.

## Acknowledgements

We thank the whole team of the Banded Mongoose Research Project that ensured the continuation of the project and were involved in collecting the invaluable life-history data and genetic samples. Sequencing was carried out by Edinburgh Genomics, The University of Edinburgh, which is partly supported with core funding from the Natural Environment Research Council (UKSBS PR18037).

## Ethical statement

Research was conducted under approval of the Uganda National Council for Science and Technology with the research registration number NS273ES, the Uganda Wildlife Authority and the corresponding reference COD/96/05 and the Ethical Review Committee of the University of Exeter. All research procedures adhered to the ASAB Guidelines for the Treatment of Animals in Behavioural Research and Teaching (ASAB Ethical Committee and ABS Animal Care Committee 2022).

