## Supplementary material for "Banded mongooses discriminate relatedness and MHC diversity in unfamiliar conspecifics"

### MATERIAL AND METHODS

*Control vs experimental presentations*

General linear mixed effect models (GLMMs) were established to test for a difference between control and experimental presentations for the different response measures. Each model included one of the six response variables: licking, marking, contact (log), sniffing (log +1), duration (log), and rolling (log +1). Some response variables were log transformed to avoid heteroscedasticity issues, and for variables with many zero values, 1 was added to include these in the log transformation. Each model also included the type of presentation (control or experimental) as an explanatory variable, and the identity of the odor recipient and the pack they reside in as random effects.

### RESULTS

*Correlational analyses*

We did not find a strong correlation ($r < 0.3$ in all cases) between microsatellite-derived measures and MHC-derived measures (Tab. S1). However, strong significant correlations ($r > 0.7$) were detected between MHC diversity measured as distinct alleles and distinct supertypes as well as MHC similarity of alleles and supertypes (Tab. S1). Nonetheless these MHC measures were used for further investigation, as they were not fitted simultaneously in a model and they contain information on MHC functional diversity on different scales.

We did not find strong correlations between the three behavioral response variables that we included in our GLMMs; Contact, Sniffing and Duration ($r < 0.3$ in all cases) with the exception of Contact and Duration ($r = 0.698$, Tab. S2).

**Table S1 Correlational analysis for sMLH, relatedness and MHC diversity measures** Correlation estimated using Pearson's product-moment correlation analysis. Shown are the corresponding p-values and the 95% confidence intervals.

| *Variables investigated* | *r* | *p* | *Lower CI* | *Upper CI* |
|---|---|---|---|---|
| *sMLH & MHC allele number* | 0.109 | 0.258 | -0.080 | 0.290 |
| *sMLH & MHC supertype number* | 0.257 | 0.006 | 0.074 | 0.423 |
| *MHC allele number & MHC supertype number* | **0.883** | **<2.2$^{-16}$*** | 0.834 | 0.919 |
| *Relatedness & sMLH* | -0.028 | 0.6212 | -0.140 | 0.084 |
| *Relatedness & MHC allele similarity* | 0.180 | 0.317 | -0.174 | 0.493 |
| *Relatedness & MHC supertype similarity* | 0.298 | 0.092 | -0.050 | 0.582 |
| *MHC allele similarity & MHC supertype similarity* | **0.835** | **1.488$^{-09}$*** | 0.690 | 0.916 |

**Table S2 Correlational analysis for response measures** Correlation estimated using Pearson's product-moment correlation analysis. Shown are the corresponding p-values and the 95% confidence intervals.

| | *r* | *p* | *Lower CI* | *Upper CI* |
|---|---|---|---|---|
| *Contact & Duration* | 0.698 | **<2.2$^{-16}$*** | 0.636 | 0.751 |
| *Contact & Sniffing* | 0.180 | **0.002*** | 0.070 | 0.286 |
| *Contact & Licking* | 0.014 | 0.811 | -0.098 | 0.125 |
| *Contact & Rolling* | **0.737** | **<2.2$^{-16}$*** | 0.681 | 0.784 |
| *Contact & Licking* | 0.032 | 0.575 | -0.080 | 0.143 |
| *Duration & Sniffing* | 0.136 | **0.017** | 0.024 | 0.244 |
| *Duration & Licking* | -0.013 | 0.827 | -0.124 | 0.099 |
| *Duration & Rolling* | 0.635 | **<2.2$^{-16}$*** | 0.563 | 0.697 |
| *Duration & Marking* | 0.122 | **0.033*** | 0.010 | 0.230 |
| *Sniffing & Licking* | -0.030 | 0.599 | -0.141 | 0.082 |
| *Sniffing & Rolling* | -0.082 | 0.152 | -0192 | 0.030 |
| *Sniffing & Marking* | 0.139 | **0.015*** | 0.027 | 0.246 |
| *Licking & Rolling* | 0.027 | 0.636 | -0.085 | 0.138 |
| *Licking & Marking* | -0.070 | 0.221 | -0.180 | 0.042 |
| *Rolling & Marking* | -0.088 | 0.125 | -0.198 | 0.024 |

*Control vs experimental presentations*

Contact (estimate = 0.6861, SE = 0.1510, t-value = 4.542, p-value = $7.87^{-06}$), sniffing (estimate =0.4575, SE = 0.0884, t-value = 5.178, p-value = $3.97^{-07}$), and duration (estimate = 0.5017, SE = 0.1255, t-value = 3.997, p-value = $7.96^{-05}$) differed significantly between control and experimental presentations (Fig.S1, Tab. S3). For licking (estimate = 0.0705, SE = 0.0843, t-value = 0.835, p-value = 0.404), marking (estimate = 0.1650, SE = 0.1223, t-value = 1.348, p-value = 0.178) and rolling (estimate = -0.0518, SE = 0.4351, t-value = -0.119, p-value = 0.905) there was no significant difference between control and experimental treatments detectable, likely because these behaviors were relatively rare (Figure S1).

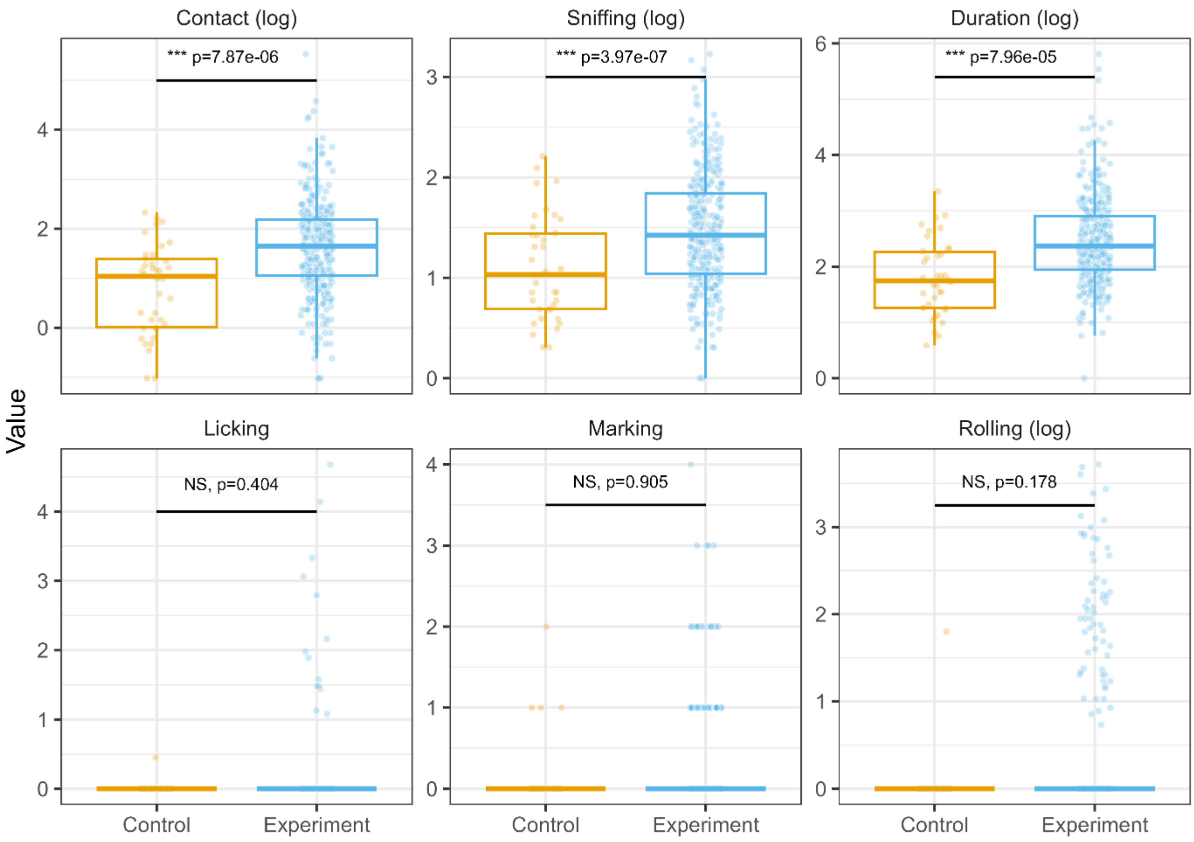


**Figure S1 Differences between control and experimental presentations.** Difference in the response values for the control and experimental presentations are shown. Boxplot whiskers show the $25^{th}$ and $75^{th}$ percentiles, the box shows the inner $50^{th}$ percentile, and the line shows the median. P-values were calculated based on Satterthwaite's method.

**Table S3 Model output for relationship between response variable and presentation type.** The table shows the model output investigating differences between control and experimental presentations using a GLMM. P-values were calculated based on Satterthwaite's method.

| *Response variable* | *estimate* | *SE* | *t-value* | *p-value* |
|---|---|---|---|---|
| *Contact* | 0.6861 | 0.1510 | 4.542 | **$7.87^{-06*}$** |
| *Sniffing* | 0.4575 | 0.0884 | 5.178 | **$3.97^{-07*}$** |
| *Duration* | 0.5017 | 0.1255 | 3.997 | **$7.96^{-05*}$** |
| *Licking* | 0.0705 | 0.0843 | 0.835 | 0.404 |
| *Rolling* | 0.1650 | 0.1223 | 1.348 | 0.178 |
| *Marking* | -0.0518 | 0.4351 | -0.119 | 0.905 |
